# Deficiency in the cell-adhesion molecule *dscaml1* impairs hypothalamic CRH neuron development and perturbs normal neuroendocrine stress axis function

**DOI:** 10.1101/2022.09.22.509087

**Authors:** Manxiu Ma, Alyssa A. Brunal, Kareem C. Clark, Carleigh Studtmann, Katelyn Stebbins, Shin-ichi Higashijima, Y. Albert Pan

**Author notes:** Correspondence should be addressed to Y. Albert Pan.

## Abstract

The corticotropin-releasing hormone (CRH)-expressing neurons in the hypothalamus are critical regulators of the neuroendocrine stress response pathway, known as the hypothalamic-pituitary-adrenal (HPA) axis. As developmental vulnerabilities of CRH neurons contribute to stress-associated neurological and behavioral dysfunctions, it is critical to identify the mechanisms underlying normal and abnormal CRH neuron development. Using zebrafish, we identified *Down syndrome cell adhesion molecule like-1 (dscaml1*) as an integral mediator of CRH neuron development and necessary for establishing normal stress axis function. In *dscaml1* mutant animals, hypothalamic CRH neurons had higher *crhb* (the CRH homolog in fish) expression, increased cell number, and reduced cell death compared to wild-type controls. Physiologically, *dscaml1* mutant animals had higher baseline stress hormone (cortisol) levels and attenuated responses to acute stressors. Together, these findings identify *dscaml1* as an essential factor for stress axis development and suggest that HPA axis dysregulation may contribute to the etiology of human *DSCAML1* -linked neuropsychiatric disorders.

## INTRODUCTION

The hypothalamic corticotropin-releasing hormone (CRH)-expressing neurons are the central regulators of the neuroendocrine stress response pathway, known as the hypothalamic-pituitary-adrenal (HPA) axis in mammals or the hypothalamic-pituitary-interrenal (HPI) axis in fish (Denver, 2009). Upon exposure to environmental disturbances (i.e., stressors), stress-related neural inputs converge on hypothalamic CRH neurons to activate a hormonal cascade that ultimately leads to the release of glucocorticoids, which broadly affects cognitive, affective, metabolic, and immune functions (Spencer and Deak, 2017; McEwen and Akil, 2020).

The development of CRH neurons has profound effects on the function of the HPA and HPI axis (collectively referred to as the stress axis). Developmental perturbations of CRH neurons, particularly in early-life periods, lead to long-term changes in CRH neuron function (Regev and Baram, 2014). Additionally, rodent models demonstrate that dysregulation of CRH neurons increases anxiety- and depressive-like phenotypes (Keen-Rhinehart et al., 2009; Kolber et al., 2010). These studies underscore the need to identify the genes and molecules mediating CRH-neuron development and determine how developmental perturbations affect stress axis function.

During early development, hypothalamic CRH neurons are generated from progenitors in the ventral diencephalon (Alvarez-Bolado, 2019; Nagpal et al., 2019; Placzek et al., 2020). Secreted factors including FGF10, SHH, BMPs, and Nodal first define the anterior-dorsal hypothalamic domain; within this domain, CRH neurons are progressively specified by a combination of key transcription factors, including Fezf2, Otp, Sim1, Arnt2, and Brn2. Once specified, CRH neurons require further differentiation, such as neuronal morphogenesis, synaptogenesis, epigenetic programming, and cell death, to establish a functional stress-responsive neural circuit. These later stages of neuronal differentiation are shaped by intercellular interactions mediated by membrane-localized cell-adhesion molecules (Moreland and Poulain, 2022). The specific cell-adhesion molecules that mediate developmental signaling in CRH neurons are still unknown.

To address this knowledge gap, we utilized zebrafish (*Danio rerio*) as the model. The structure and function of the mammalian HPA axis and the teleostean HPI axis are highly conserved (Wendelaar Bonga, 1997; Lohr and Hammerschmidt, 2011). In mammals, the CRH neurons that are involved in HPA-axis activation are in the hypothalamic paraventricular nucleus (PVN). In teleosts (ray-finned fish), the neuroendocrine preoptic area (NPO) is ontogenically equivalent to the PVN, and CRH neurons within the NPO perform similar roles as their mammalian counterparts (Herget and Ryu, 2015). A unique advantage of the zebrafish system is its rapid and external development. The development of the HPI axis begins between 1-2 days post fertilization (dpf), and stress-induced cortisol signaling and behaviors can be observed at 4-5 dpf (Alsop and Vijayan, 2008; Alderman and Bernier, 2009; Bai et al., 2016). The rapid development and translucency of zebrafish allow direct microscopic observation of CRH neuron development in intact, developing animals.

In this study, we investigated the roles of a conserved neuronal signaling molecule— *Down syndrome cell adhesion molecule-like 1* (zebrafish gene: *dscaml1;* mouse gene: *Dscaml1;* Human gene: *DSCAML1;* Protein: DSCAML1). DSCAML1 is one of two DSCAM family members in vertebrates, the other being DSCAM (Garrett et al., 2012). Unlike invertebrate DSCAMs, vertebrate DSCAMs are not significantly alternatively spliced (Sanes and Zipursky, 2020). In the mammalian retina, DSCAML1 prevents excessive aggregation between cells and promotes developmental cell death (Fuerst et al., 2009; Garrett et al., 2016). DSCAML1 also acts to refine synaptic specificity and synapse number (Yamagata and Sanes, 2008; Sachse et al., 2019). In humans, rare variants in *DSCAML1* are associated with several neurodevelopmental disorders, including autism spectrum disorder, cortical abnormality, and epilepsy (Iossifov et al., 2014; Karaca et al., 2015; Hayase et al., 2020; Ogata et al., 2021). Genetic and epigenetic studies also implicate *DSCAML1* in the stress response to violent experiences (Caramillo et al., 2015; Saadatmand et al., 2021). However, the relationship between *DSCAML1* and the stress axis remains unknown.

Using zebrafish, we previously explored how DSCAML1 affects neural pathways and systemic functions (Ma et al., 2020a; Ma et al., 2020b). We found that *dscaml1* deficiency resulted in various physiological and behavioral deficits, including darker pigmentation, slower light adaptation, and slower eye movements (saccades) (Ma et al., 2020b). Interestingly, darker pigmentation and slower light adaptation can also be caused by abnormal glucocorticoid receptor signaling, as seen in the zebrafish *glucocorticoid receptor* (*gr*) mutants, suggesting that the stress axis may be dysfunctional in *dscaml1* mutants (Griffiths et al., 2012; Muto et al., 2013).

Here, we report that *dscaml1* deficiency in zebrafish perturbs CRH neuron development and impairs the normal function of the HPI axis. These findings show that DSCAML1 is necessary for stress axis development and raise the possibility that stress dysfunction contributes to human *DSCAML1-linked* disorders.

## RESULTS

### *dscaml1* deficiency results in overexpression of neuroendocrine factors

To gain an unbiased view of the molecular changes resulting from *dscaml1* deficiency, we compared the transcriptomic profiles between *dscaml1* homozygous mutant (*dscaml1-/-*) and control (wild type) animals using RNA sequencing (RNA-seq). cDNA from whole 3.5-4 days post-fertilization (dpf) *dscaml1-/-* and control larvae were sequenced using an Illumina next-generation sequencer. Using a threshold of at least two-fold change and an adjusted *p-* value of less than 0.01, we identified 25 upregulated and 79 downregulated genes (Fig. 1A, Supplementary file 1).

**Fig. 1.**
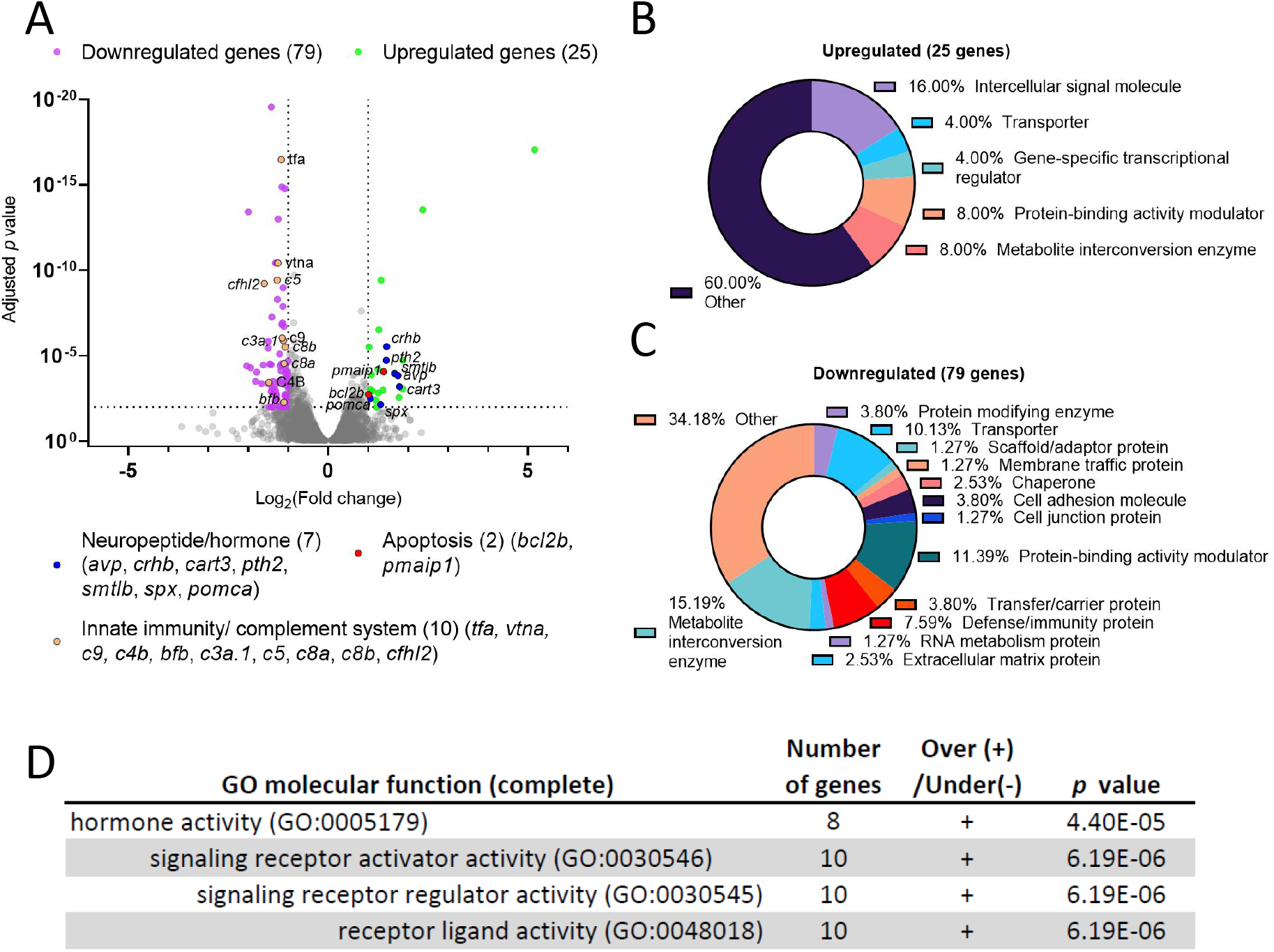
Differential gene expression analysis of *dscaml1* deficient zebrafish. (A) Volcano plot of relative gene expression in *dscaml1-/-* versus control animals. Each dot represents an individual gene, with colored dots representing gene groups as indicated on the graph. The dotted lines show the significance level (adjusted *p* < 0.01) and fold change (increase or decrease by two-fold or more) thresholds. (B, C) Protein class categorization analysis for upregulated (B) and downregulated (C) genes. (D) Table of significantly enriched (*p<0.05*) GO terms for molecular function.

Among the 25 upregulated genes in the *dscaml1-/-* animals, 7 were secreted neuropeptides/hormones expressed in the hypothalamus or pituitary: *corticotropin-releasing hormone b (crhb), parathyroid hormone 2 (pth2), somatolactin beta (smtlb), cocaine- and amphetamine-regulated transcript 3 (cart3), proopiomelanocortin a (pomca), arginine vasopressin (avp*), and *spexin hormone (spx*). Among them, three (*crhb, avp, pomca*) are core regulators of the HPI axis (Alsop and Vijayan, 2009; Lohr and Hammerschmidt, 2011). *crhb* encodes the zebrafish CRH; *avp* encodes the neuropeptide AVP that controls osmolarity, blood pressure, and synergizes with CRH to promote cortisol release; *pomca* encodes the adrenocorticotropic hormone (ACTH), the primary pituitary hormone that triggers glucocorticoid release. Additionally, two genes involved in apoptosis are upregulated: *BCL2 apoptosis regulator b (bcl2b*) and *phorbol-12-myristate-13-acetate-induced protein 1 (pmaip1/noxa*). Protein class categorization analysis with PANTHER revealed “intercellular signal molecule” as the largest class (4 genes, 16%) (Fig. 1B) (Mi et al., 2021).

The 79 genes downregulated in the *dscaml1-/-* animals are more diverse in function compared to the upregulated genes. Protein class categorization analysis with PANTHER identified the largest protein classes as metabolite interconversion enzymes (12 genes, 15.19%), protein-binding activity modulators (9 genes, 11.39%), transporters (8 genes, 10.13%), and defense/immunity protein (6 genes, 7.59%) (Fig. 1C). We noted that 30 of the 79 (38%) downregulated genes are highly expressed in the liver, and 10 of these genes are involved in innate immunity and the complement cascade (Fig. 1A, Supplementary file 2). These results suggest that liver function and innate immunity may be suppressed in the *dscaml1* mutants.

To identify the signaling pathways affected by *dscaml1* deficiency, we analyzed all differentially expressed genes (*p<0.01*, 210 mapped genes) using the statistical enrichment test (PANTHER Classification System, version 17.0) (Mi et al., 2021). All significantly enriched (FDR*<0.05*) Gene Ontogeny (GO) terms for molecular function relate to the parent GO term *hormone activity* (Fig. 1D). The PANTHER pathway analysis also identified another significant stress-modulating neuropeptide, *adenylate cyclase activating polypeptide 1b (adcyap1b*, also known as *PACAP*) that is significantly upregulated (0.66 fold change, adjusted *p<0.0001*) (Stroth et al., 2011).

Together, our transcriptomic analyses indicate that *dscaml1* deficiency results in the upregulation of neuropeptide/hormonal signaling and the downregulation of liver and innate immune function. Suppression of liver and immune function is a hallmark of stress axis activation, which is consistent with the overexpression of the principal neuropeptides involved in the stress axis (*crhb, avp, pomca*, and *adcyap1b*).

### *dscaml1* deficiency alters the development of CRH neurons in the NPO

To further investigate whether the stress axis is perturbed in *dscaml1* mutants, we examined the development of CRH neurons in the NPO (CRH^NPO^ neurons), which are characterized by their expression of *crhb* (Fig. 2A) (Herget et al., 2014; Vom Berg-Maurer et al., 2016). We focused on three developmental stages (2, 3, and 5 dpf) that span the period between the first appearance of *crhb+* neurons in the NPO (2 dpf) and the onset of stress axis responsivity (4-5 dpf) (Chandrasekar et al., 2007; Alderman and Bernier, 2009; Clark et al., 2011).

**Fig. 2.**
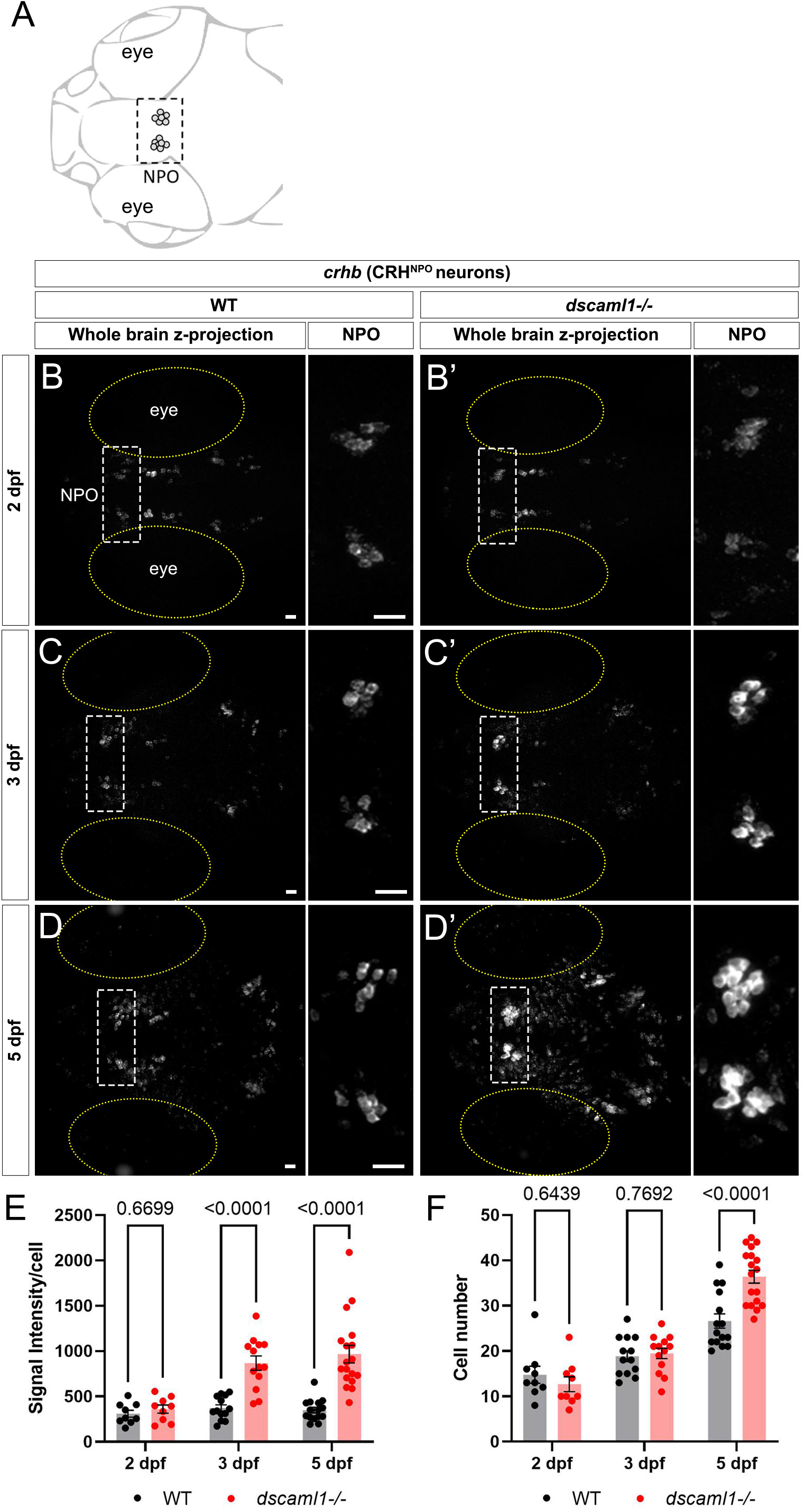
*dscaml1* deficiency alters the development of CRH^NPO^ neurons. (A) Illustration of CRH neurons (gray circles) in the NPO (boxed area) in the larval zebrafish. (B-D’) Developmental trajectory of CRH^NPO^ neurons, labeled by *crhb* FISH. At each developmental stage, a representative confocal z-stack projection of the whole brain is shown in the left panel, and the substack containing the NPO is enlarged and shown in the right panel. The white boxes indicate the locations of the NPO, and the yellow ovals mark the eyes. Wild-type (WT) animals are shown in panels B, C, and D. *dscaml1-/-* animals are shown in panels B’, C’, and D’. (E-F) Quantification of the signal intensity per cell (E) and cell number (F). Multiple-comparison corrected *p* values are as shown. WT: n=9 (2 dpf), 13 (3 dpf), 15(5 dpf). *dscaml1-/-:* n=9 (2 dpf), 13 (3 dpf), 18 (5 dpf). Scale bars are 20 μm. Mean, standard error, and corrected *p* values are shown.

Using fluorescent *in situ* hybridization (FISH), we found that *crhb* expression pattern was initially similar between *dscaml1-/-* and wild-type (WT) at 2 dpf (Fig. 2B-B’). At 3 dpf, we began to see higher *crhb* expression in the NPO of *dscaml1-/-* animals (Fig. 2C-C’). At 5 dpf, there is widespread overexpression of *crhb* in *dscaml1-/-* animals, with *crhb* FISH intensity most notably elevated in the NPO (Fig. 2D-D’). Quantification of *crhb* FISH signal intensity among CRH^NPO^ neurons showed that *crhb* expression is higher in *dscaml1-/-* animals, compared to wild-type animals (WT) (Fig. 2E). There was a significant difference in signal intensity per cell by developmental stage and genotype, and a significant interaction between stage and genotype (two-way ANOVA, Supplementary Table I). Pair-wise comparisons with Holm-Sidak correction found a significant increase at 3 and 5 dpf but not at 2 dpf (Fig 2E).

In addition to changes in *crhb* expression levels, *dscaml1* deficiency also increased the number of *crhb*-expressing CRH^NPO^ neurons. In WT animals, the number of CRH^NPO^ neurons increased over time, from 14.78 cells (2 dpf) to 18.85 cells (3 dpf) to 26.60 cells (5 dpf) per animal (Fig 2B, C, D, F). In *dscaml1-/-* animals, the number of CRH^NPO^ neurons increased at a higher rate, from 12.67 cells (2 dpf) to 19.46 cells (3 dpf) to 36.39 cells (5 dpf) per animal (Fig. 2B’, C’, D’, F). There was a significant difference in cell number by developmental stage and genotype, and there was a significant interaction between stage and genotype (Two-way ANOVA, Supplementary Table I). Pair-wise comparisons with Holm-Sidak correction found a significant increase in cell number between *dscaml1-/-* and WT animals at 5 dpf but not at 2 and 3 dpf (Fig 2F, adjusted *p* values as shown).

Overall, we found significant increases in *crhb* expression and cell number in CRH^NPO^ neurons in *dscaml1-/-* mutants as compared to WT. These phenotypes were not due to the visual deficits in *dscaml1-/-* animals (Ma et al., 2020b), as similar phenotypes were observed in animals raised in the dark (Supplementary Fig. 1). Together, these findings show that *dscaml1* is essential for the normal developmental trajectory of CRH^NPO^ neurons.

### *dscaml1* is essential for normal CRH^NPO^ neuron cell death

Based on the finding that mouse DSCAML1 promotes programmed cell death (PCD) in the retina (Garrett et al., 2016) and our transcriptomic analysis that showed *dscaml1* mutants expressing higher levels of genes involved in regulating apoptosis (Fig. 1A), we hypothesized that *dscaml1* deficiency might impair normal CRH^NPO^ neuron cell death. To test this hypothesis, we tracked the fate of individual CRH^NPO^ neurons using *in vivo* time-lapse imaging at 3-5 dpf, when cell number begins to diverge between *dscaml1-/-* and WT animals.

To visualize CRH^NPO^ neurons in live zebrafish, we generated a *crhb* knock-in fluorescent reporter line using CRISPR-mediated genomic insertion (Fig. 3A) (Kimura et al., 2014). A Cre-switchable *hsp-LoxP-RFP-LoxP-GFP* cassette was inserted 35 base pairs upstream of the first exon of *crhb* so that the expression of RFP (default) or GFP (with Cre-mediate recombination) would mark the endogenous *crhb*-expressing cells (Fig. 3B). The resulting transgenic line, *crhb:LoxP-RFP-LoxP-GFP (crhb:LRLG*), has RFP and GFP expression pattern that matches the endogenous *crhb* transcript expression patterns (Fig. 3C-D, compare to Fig. 2D). To validate the fidelity of fluorescent reporter expression, we examined whether *crhb:LRLG-labeled* cells in the NPO express CRH protein. In animals not exposed to Cre (default RFP expression), we found that most RFP+ neurons are CRH immunopositive (84.71±2.19%, Supplementary Fig. 2A-B). Together, these results indicate that the *crhb:LRLG* line reliably labels CRH^NPO^ neurons.

**Fig. 3.**
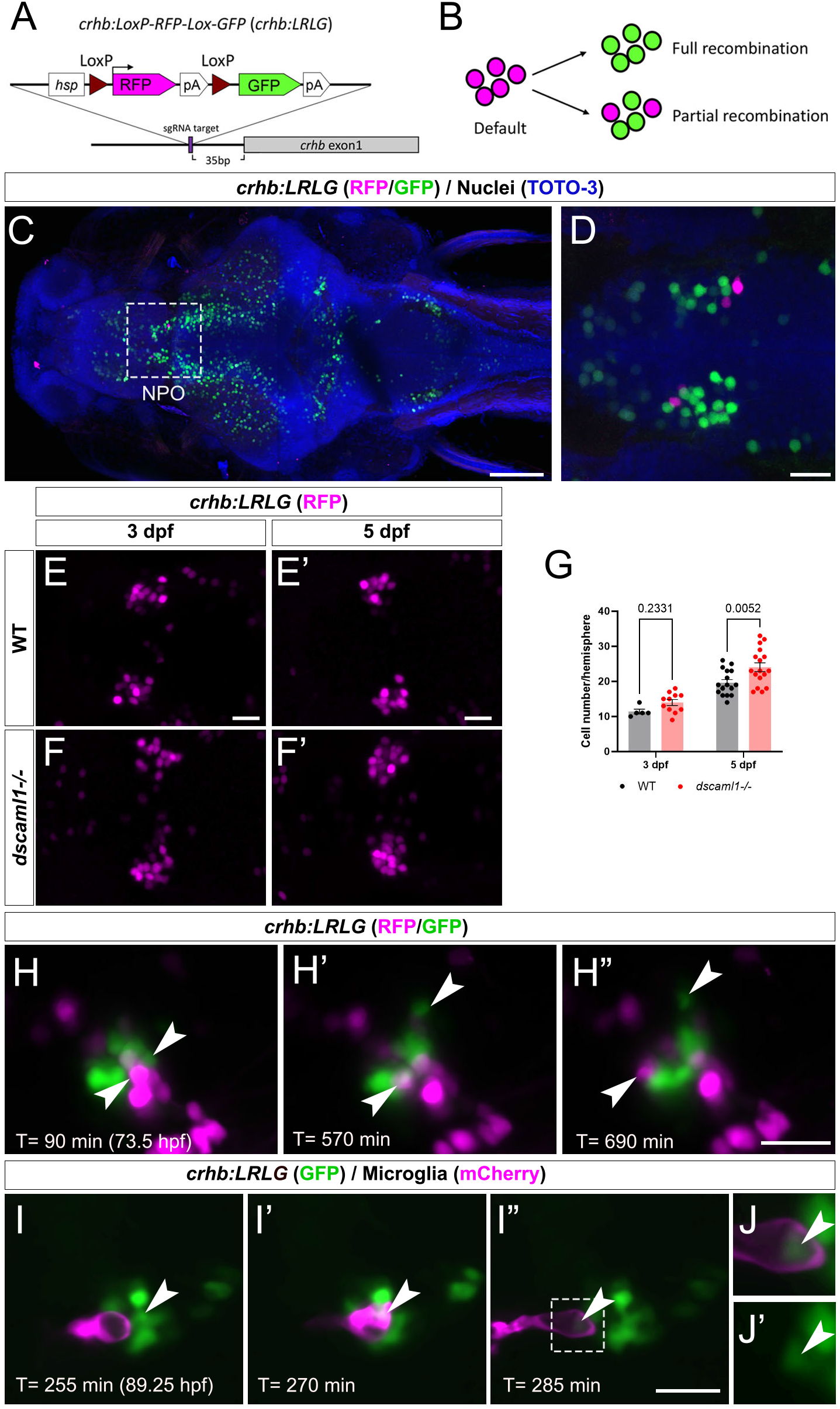
Fluorescent labeling and live imaging of CRH^NPO^ neurons. (A) Schematic of CRISPR -mediated knock-in of the *hsp-Lox-RFP-Lox-GFP* cassette at the sgRNA target site, located 35 bp upstream of exon 1 of *crhb*. The orientation and junctional structure of insertion have not been determined. (B) Schematic of *crhb:LRLG* expression. Each circle represents a fluorescent cell. Without Cre (default), RFP is expressed in all cells. With full recombination, all cells express GFP. Partial recombination results in mosaic RFP and GFP labeling. (C) Dorsal view of a fixed 5 dpf *crhb:LRLG* larvae with partial recombination stained with anti-RFP (magenta) and anti-GFP (green). The boxed area marks the NPO. (D) Higher magnification image of the NPO. Both RFP and GFP-positive neurons can be seen. (E-F’) Images of anti-RFP stained *crh:LRLG* animals without recombination. Representative wildtype (WT, E-E’) and *dscaml1-/-* (F-F’) NPO neurons are shown. (G) Quantification of RFP-positive cells at 3 and 5 dpf. WT: n=5 (3 dpf), 16 (5 dpf). *dscaml1-/-*: n=11 (3 dpf), 17 (5 dpf). Mean, standard error, and corrected *p* values are shown. (H-H”) Live *crhb:LRLG* larvae with partial recombination were imaged from 72 to 84 hpf. Three time points are shown here. Two cells (arrowheads, one green and one magenta) move away over time. (I-I”) Live *crhb:LRLG;mpeg1: Gal4;UAS:NTR-mCherry* larvae with CRH neurons labeled with GFP (green) and microglia labeled with mCherry (magenta). In this image series, one CRH neuron (arrowhead) is engulfed (I’) and then removed (I”) by a microglial cell. Images are confocal optical sections. (J-J’) Panels showing enlarged views of the boxed area in I”, with (J) or without (J’) the mCherry channel. The remnant of the CRH neuron can still be seen inside the microglia (arrowheads). Scale bars are 100 μm (panel C) or 20 μm (all other images).

In *crhb:LRLG* animals, *dscaml1* deficiency increased the number of RFP-labeled CRH^NPO^ neurons (Fig. 3E-F’). There were significant differences by age and genotype, with no significant interaction (two-way ANOVA, Supplementary Table I). There was a significantly higher number of RFP-positive neurons in the *dscaml1* mutants at 5 dpf but not 3 dpf (Fig. 3G, multiple comparison test with Holm-Sidak correction). This result corroborates our *crhb* FISH results that show increased CRH^NPO^ neuron number in *dscaml1* mutants (Fig. 2F).

Using the *crhb:LRLG* line, we first performed time-lapse imaging in anesthetized animals. We induced partial Cre-mediated recombination by injecting *CreER* mRNA into *crhb:LRLG* animals at the 1-cell stage and adding 4-hydroxytamoxifen (4-OHT) to activate CreER at 6-24 hpf (Fig. 3B). At 3-4 dpf, intermingled GFP+ and RFP+ cells can be seen in the NPO by confocal imaging (Fig. 3H-H” and Supplementary video 1). Over time, some labeled cells moved away from the CRH^NPO^ neuron cluster. These cells were likely dying cells being carried away by microglia, as seen in previous zebrafish studies (Mazaheri et al., 2014). To confirm this, we induced GFP expression (by injecting codon-optimized *Cre* mRNA)(Horstick et al., 2015) in all *crhb:LRLG-labeled* cells and labeled microglia with the *mpeg1:mCherry* transgene (Fig. 3I-J” and Supplementary video 2) (Espenschied et al., 2019). Indeed, mCherry+ microglia migrated toward the GFP+ cell cluster, engulfed GFP-positive CRH^NPO^ neurons, and carried the engulfed cells away. The remnant of the engulfed cell can be seen inside a large vacuole within the microglia (Fig. 3J-J’). These results suggest that CRH^NPO^ neurons undergo PCD and that dying cells are rapidly removed by microglia.

Next, to track the fate of individual CRH^NPO^ neurons, we performed two-photon imaging from 3 to 5 dpf on *crhb:LRLG* animals with partial Cre-mediated recombination (Fig. 4A). To minimize the potential effects of stress on PCD (Irles et al., 2014), we anesthetized and immobilized the animals during imaging. In between imaging sessions, each animal is allowed to recover in individual wells of 12-well plates, under normal light-dark cycles. Cells that are present at the first time point (78 hpf) were tracked at four subsequent time points (84, 96, 108, and 120 hpf) and categorized as either persisting (present at the last time point) or lost (lost at any of the following time points) (Fig. 4B-B”). Overall, we observe a trend of reduced cell loss in the *dscaml1*+/- and *dscaml1-/-* animals (Chi-square test for trend, *p*=0.0398) (Fig. 4C, total number of tracked cells as indicated). In WT animals, 16.41% of cells are lost, versus 10.91% and 7.53% for *dscaml1+/- and dscaml1-/-*, respectively.

**Fig. 4.**
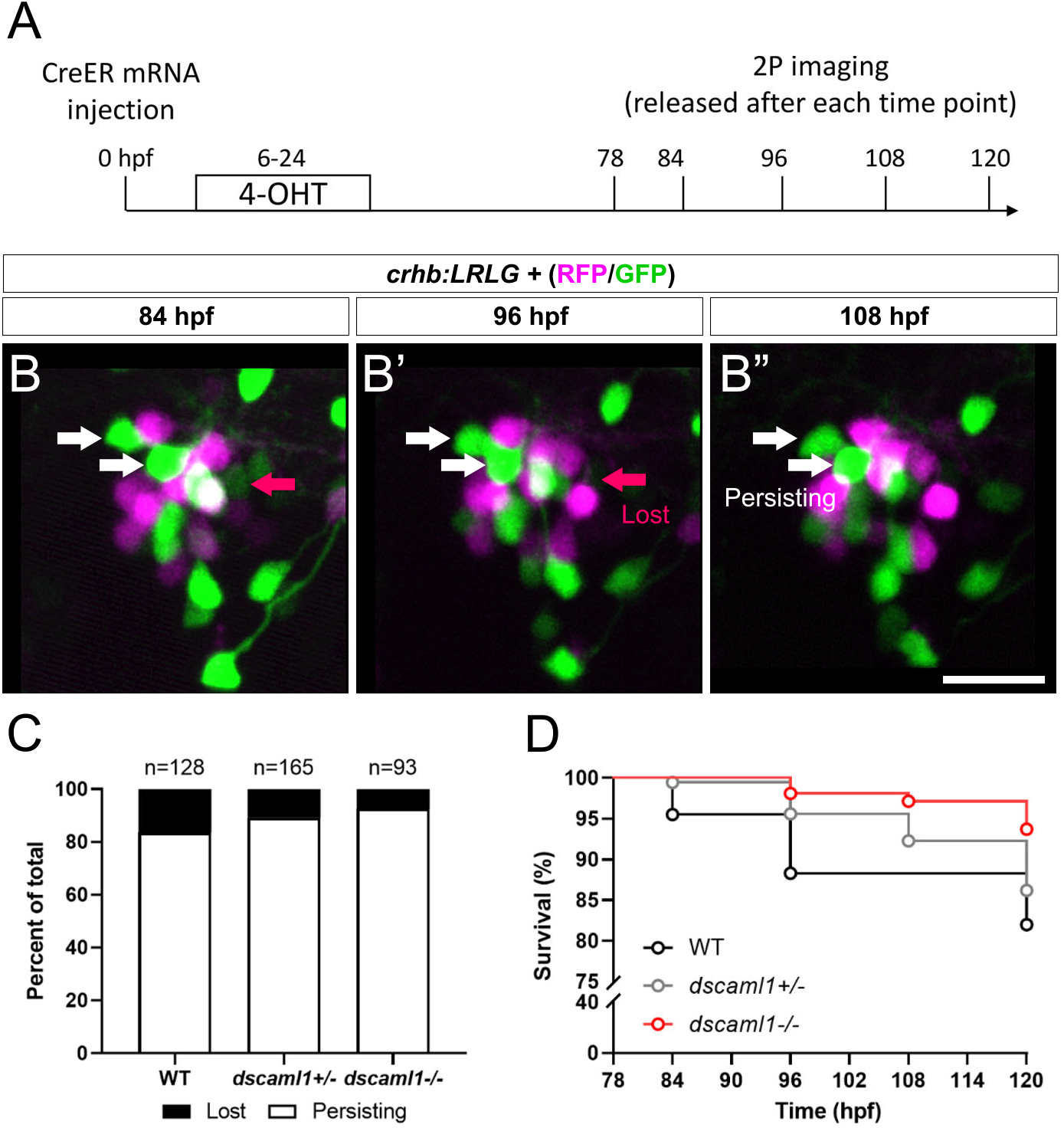
Live tracking of CRH^NPO^ neuron cell fate. (A) Timeline of time-lapse two-photon (2P) imaging experiment. Partial recombination of *crhb:LRLG* was induced by 4-OHT at 6-24 hpf, and imaging was performed at 78, 84, 96, 108, and 120 hpf. Animals were briefly anesthetized during imaging and allowed to recover in between imaging sessions. (B-B”) Tracking of individual CRH^NPO^ neurons. Three example time frames are shown. Individual fluorescent cells can be tracked over time and are divided into two categories: persisting (white arrows) or lost (pink arrow). (C) Quantification of the percentage of persisting versus lost CRH^NPO^ neurons. Sample size (cell number) as indicated for each genotype. (D) Survival curve of individual CRH^NPO^ neurons in each genotypic group.

Finally, considering the timing of cell loss, we plotted the survival curve of CRH^NPO^ neurons for each genotype. Again, there was a significant difference in the trends of cell loss (Logrank test for trend, *p=*0.0098), with the *dscaml1-/-* animals consistently showing a higher survival rate (Fig. 4D). Together, these results show that *dscaml1* deficiency reduces PCD of CRH^NPO^ neurons.

### Stress axis function is perturbed in *dscaml1* mutant animals

Given the developmental perturbation of CRH^NPO^ neurons as well as the global changes in gene expression related to stress axis activation, we next determined whether the hormonal output of the stress axis—cortisol—is altered. Cortisol levels were measured using an enzyme-linked immunosorbent assay (ELISA) on homogenates made from pools of 5 dpf animals (30 animals per sample, 6 samples per condition). All animals were raised under standardized conditions, at the same density, and with a normal circadian cycle (14 h day/10 h night) (Yeh et al., 2013). Under this circadian cycle, *dscaml1* mutant animals exhibit similar diurnal locomotor rhythms as wild-type animals (Ma et al., 2020b).

Baseline and stressed conditions were tested to evaluate potential alterations of cortisol in control (WT and *dscaml1 +/-*) and *dscaml1-/-* animals at 5 dpf. To measure baseline cortisol, we collected unperturbed animals within 30 min after light onset (zeitgeber time 0, ZT0) and in the afternoon (ZT6-8). To measure stress-induced cortisol, animals were exposed to either stirring stress (swirling water, 5 minutes) (Castillo-Ramirez et al., 2019) or hyperosmotic stress (250 mM NaCl, 20 minutes) at ZT6-8 (Yeh et al., 2013). These acute stressors are well-characterized and are comparable to the water current and salinity changes experienced by zebrafish larvae in their natural habitat (Clark et al., 2011).

At baseline, *dscaml1-/-* animals had significantly higher cortisol levels than control animals (Multiple Mann-Whitney test with Holm-Sidak correction. Adjusted *p* values shown in Fig. 5A). At ZT0, the *dscaml1* mutant baseline cortisol levels were 2.7-fold higher than that of controls (median 350.06 pg/ml versus 129.80 pg/ml). The difference in cortisol was less pronounced at ZT6-8, but *dscaml1* mutants still had 2-fold higher cortisol levels than controls at baseline (median 238.508 pg/ml versus 118.516 pg/ml).

**Fig. 5.**
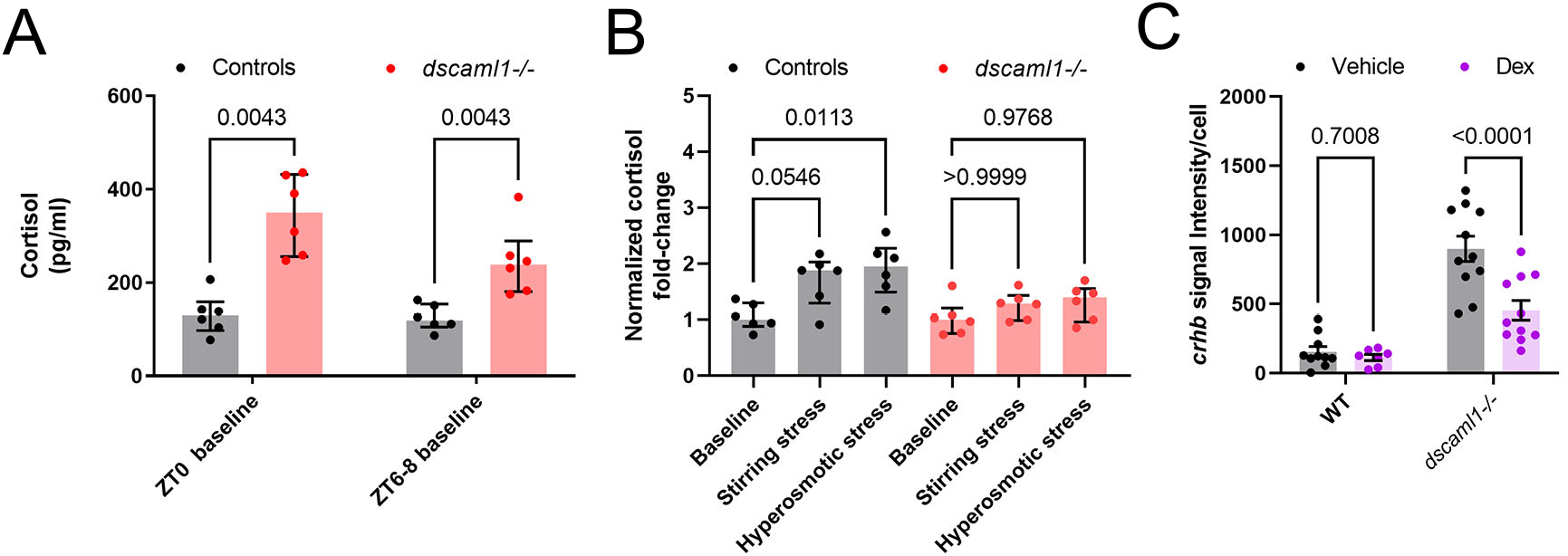
Cortisol levels and response to exogenous glucocorticoids. (A-B) Cortisol profile for 5 dpf larvae. For each sample (dot), cortisol was extracted from a pool of 30 animals. n=6 for all groups. Median, interquartile range, and corrected *p* values are shown. (A) Baseline cortisol in control (black) and *dscaml1-/-* (red) animals. (B) Baseline normalized cortisol fold change in control (black) and *dscaml1-/-* (red) animals. (C) Quantification of *crhb* signal intensity per cell in CRH^NPO^ neurons. Vehicle: n=10 (WT), 11 (*dscaml1-/-*). Dex: n=7 (WT), 11 (*dscaml1-/-*). Mean, standard error, and corrected *p* values are shown.

After acute exposure to stressors, we found that *dscaml1* mutant animals exhibited attenuated cortisol induction. We assessed the extent of stress-induced cortisol production by normalizing cortisol levels to the baseline cortisol of the same genotype at the same circadian time (ZT6-8). In the control group, stirring stress and hyperosmotic stress produced 1.88 and 1.95-fold increases in cortisol over the control baseline, respectively (grey bars, Fig. 5B). The response to hyperosmotic stress was more robust (*p*=0.0113, Multiple Mann-Whitney tests with Holm-Sidak correction) than that generated by stirring stress (*p*=0.0546). In the *dscaml1-/-* group, stirring stress and hyperosmotic stress only produced 1.29 and 1.4-fold increases in cortisol over the *dscaml1-/-* baseline, respectively (red bars, Fig. 5B). These increases were not statistically significant (*p*>0.9999 for stirring stress, *p*=0.9768 for hyperosmotic stress).

Together, these results show that *dscaml1* deficiency elevates baseline cortisol levels and imparing responses to acute stressors. These findings indicate that *dscaml1* is critical for establishing the normal function of the stress axis.

### *dscaml1* mutants are responsive to glucocorticoids

Some aspects of the stress axis-related phenotypes in *dscaml1* mutants resemble the zebrafish *glucocorticoid receptor (gr*) mutants. In particular, both *gr* and *dscaml1* mutants exhibit elevated baseline cortisol and *crhb* (Ziv et al., 2013). This resemblance raises the possibility that the *dscaml1* mutant phenotypes may result from insufficient glucocorticoid receptor-mediated signaling. To test this, we examined whether *dscaml1* mutants can respond transcriptionally to exogenously applied glucocorticoids. We examined the expression of *crhb* in the NPO, as it is under feedback control from glucocorticoid-glucocorticoid receptor signaling (Watts, 2005).

A synthetic glucocorticoid receptor agonist, dexamethasone (Dex), was added to the embryo media at a final concentration of 2 μM from 4 to 5 dpf (24 hours). Vehicle (0.02% ethanol) treated siblings were used for comparison. For *crhb* transcript level in CRH^NPO^ neurons, we found significant differences caused by Dex treatment and genotype, with significant interaction between the two (two-way ANOVA, Supplementary Table I). Multiple comparison tests found that Dex significantly reduced *crhb* transcript level in *dscaml1-/-* animals but not in wild-type animals (Holm-Sidak correction, Fig. 5C with *p* values as indicated). These results show that *dscaml1* deficiency does not result in a loss of glucocorticoid responsivity in CRH^NPO^ neurons. Instead, *dscaml1* deficiency may render animals more sensitive to the inhibitory effects of glucocorticoids.

## DISCUSSION

The present study provides evidence that *dscaml1* regulates the development of hypothalamic CRH neurons and is necessary for normal stress axis function. At the transcriptome level, *dscaml1* deficiency in zebrafish results in gene expression changes that suggest stress axis hyperactivation. At the cellular level, we find that *dscaml1-/-* hypothalamic CRH neurons (CRH^NPO^ neurons) have increased stress axis-associated neuropeptide (*crhb*) expression, increased cell number, and reduced cell death. Physiologically, *dscaml1* deficiency impairs normal neuroendocrine stress axis function, which is potentially caused by developmental deficits in CRH^NPO^ neurons as well as systemic changes in hormone/neuropeptide signaling (Fig. 6). Together, these findings link DSCAML1 to the development of the stress axis and shed new light on the potential etiology of human *DSCAML1-linked* mental health conditions.

**Fig. 6.**
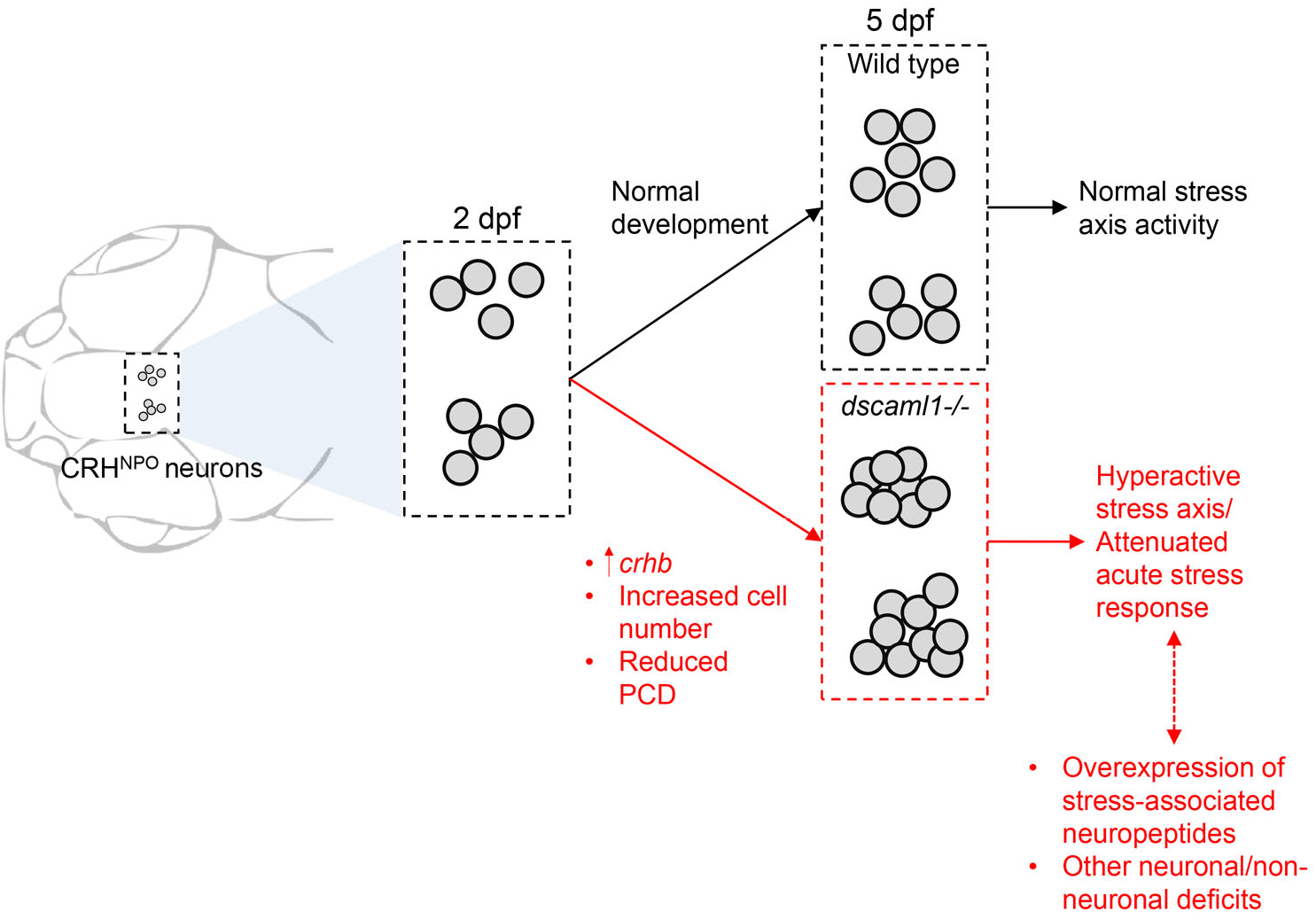
Summary of findings. Schematic summary of findings. CRH^NPO^ neurons are formed normally early on (2 dpf) but begin to exhibit developmental abnormalities in *dscaml1* deficient animals. The developmental deficits of CRH^NPO^ neurons and other systemic developmental deficits likely contribute collectively to cause the hyperactive stress axis and attenuated acute stress response in *dscaml1* mutants.

### DSCAML1 is a novel intercellular signaling molecule for CRH neuron development

A major finding of this study is that DSCAML1 is necessary for the development of hypothalamic CRH neurons. To our knowledge, DSCAML1 is the first intercellular signaling molecule to be implicated in CRH neuron development. Our finding also provides the first link between DSCAML1 and hypothalamus development. We show that, similar to retinal neurons, the regulation of cell number by PCD is a DSCAML1-mediated process in CRH neurons (Garrett et al., 2016). Further investigations are required to determine whether other aspects of DSCAML1 function, such as the regulation of cellular spacing and synaptogenesis, are involved in CRH^NPO^ neuron development (Fuerst et al., 2009; Yamagata and Sanes, 2010; Sachse et al., 2019). Additionally, given that *dscaml1* is expressed broadly in the nervous system (Ma et al., 2020b), including CRH neurons and non-CRH neurons (Supplementary Figure 3), it will be important to address whether *DSCAML1* acts cell-autonomously in CRH^NPO^ neurons.

### Regulation of CRH neuron cell death by DSCAML1

During development, PCD is critical for eliminating transient cell types, matching input and output cell populations, and maintaining cellular spacing (Yamaguchi and Miura, 2015; Wong and Marin, 2019). It has been hypothesized that PCD in the hypothalamus may specify neural circuit assembly and shape output activity (Simerly, 2002; Forger, 2009). Congruent with this idea, reduced hypothalamic PCD caused by early-life stress is associated with increased sensitivity to acute stressors in adulthood (Zhang et al., 2012; Irles et al., 2014).

As an intercellular signaling molecule, DSCAML1 may act to transduce extracellular cell death cues. One potential cue is synaptic activity, which is critical for neuronal survival during development (Wong and Marin, 2019). In culture, DSCAML1 is localized to excitatory synapses (Yamagata and Sanes, 2010) and inhibits excitatory synaptogenesis when overexpressed (Sachse et al., 2019). It remains to be determined whether DSCAML1 deficiency increases excitatory synaptic transmission and activates activity-dependent cell survival pathways in CRH neurons (Wong and Marin, 2019).

### Stress axis dysfunction in DSCAML1 deficient animals

*dscaml1-/-* animals exhibit multiple signs of stress axis activation at baseline, including cortisol elevation and the suppression of immunity-associated genes. The elevated baseline cortisol levels likely resulted in attenuated responses to acute stressors, similar to animals under chronic cortisol administration (Barton et al., 1987; Johnson et al., 2006). A likely cause of these phenotypes is the overexpression of *crhb*. In mice, broad overexpression of CRH leads to elevated corticosterone (the stress glucocorticoid in rodents) and produces phenotypes similar to Cushing’s syndrome, a human disorder caused by the overproduction of cortisol (Stenzel-Poore et al., 1992; Arnett et al., 2016).

Beyond dysfunction of hypothalamic neurons and CRH signaling, a broader neurological imbalance can also activate the stress axis. For example, seizures have been shown to activate the HPA axis, which increases the likelihood of future seizures (O’Toole et al., 2014; Hooper et al., 2018). It has been reported that *Dscaml1* mutant rats and human patients with *DSCAML1* loss-of-function variants exhibit neuronal hyperactivation and seizures (Hayase et al., 2020). It is, therefore, possible that excitation-inhibition imbalance in extra-hypothalamic regions may result in increased CRH^NPO^ neuron firing in zebrafish *dscaml1* mutants. Further studies on the cell-type-specific functions of *dscaml1* are needed to understand the precise cause of stress axis hyperactivation in *dscaml1-/-* animals.

### The interplay between cortisol signaling and CRH neuron development

Facilitation and feedback are signature features of the stress axis (Dallman et al., 1992; Spencer and Deak, 2017). While CRH neurons control cortisol levels, cortisol signaling also affects the development of CRH neurons. Zebrafish *gr* mutants have phenotypes similar to that of *dscaml1* -/- animals, including slow visual-background adaptation, sluggish light onset response, elevated cortisol, and increased expression of *crhb* and *pomca* (Griffiths et al., 2012; Muto et al., 2013; Ziv et al., 2013). Surprisingly, rather than decreased glucocorticoid receptor signaling (as in*gr* mutants), *dscaml1-/-* animals have intact glucocorticoid receptor signaling, with dexamethasone exerting strong suppression of *crhb* expression in the NPO. Thus, despite the superficial phenotypic similarity, the underlying signaling mechanisms are distinct between *dscaml1* and *gr* mutants. Nevertheless, further work is needed to disambiguate the relationship between cortisol disturbances and developmental deficits. A possible approach would be to normalize cortisol levels by genetically ablating interrenal cells (i.e., genetic adrenalectomy) and supplementing with constant levels of exogenous cortisol (Gutierrez-Triana et al., 2015).

### Conclusions

In conclusion, this work shows that DSCAML1 is integral for developing the hypothalamic neurons that regulate the neuroendocrine stress axis. Using zebrafish as a vertebrate model for the ontogenesis of the stress axis, we found that *dscaml1* deficiency results in CRH neuron deficits and dysfunction of the stress axis. Genetic perturbations of *DSCAML1* are seen in patients suffering from a wide range of mental health disorders, including intellectual disability, autism spectrum disorder, schizophrenia, epilepsy, and stress disorder (Iossifov et al., 2014; Caramillo et al., 2015; Karaca et al., 2015; Hayase et al., 2020; Ogata et al., 2021; Saadatmand et al., 2021). Developmental deficits in the stress axis may contribute to the etiology of these disorders.

## MATERIALS AND METHODS

### Zebrafish Husbandry

Zebrafish (all ages) were raised under a 14/10 light/dark cycle at 28.5°C. Embryos and larvae were raised in E3 buffer (5 mM NaCl, 0.17 mM KCl, 0.33 mM CaCl_2_, 0.33 mM MgSO_4_) (Nüsslein-Volhard et al., 2002). All zebrafish used in this study were in a mixed background of AB and TL wild-type strains (Zebrafish International Resource Center). Sex was not a relevant variable for the stages used in this study (0-6 dpf), as laboratory zebrafish remain sexually undifferentiated until two weeks of age (Maack and Segner, 2003; Wilson et al., 2014). All procedures were performed according to protocols approved by the Institutional Animal Care and Use Committee at Virginia Tech and the National Institute for Basic Biology.

### Transgenic and mutant zebrafish lines

The *dscaml1^vt1^* loss-of-function allele contains a 7 base pair deletion that results in premature translational termination (Ma et al., 2020b). Animals used for live imaging were in homozygous *nacre (mitfa*) mutant background to prevent pigment formation (Lister et al., 1999). The microglia RFP line *[Tg(mpeg1:Gal4;UAS:NTR-mCherry)]* was obtained from Dr. John Rawls at Duke University (Espenschied et al., 2019). The *crhb:LoxP-RFP-LoxP-GFP* line was generated using CRISPR-mediated knock-in, as described by Kimura et al. (Kimura et al., 2014). The sgRNA sequence for the *crhb* knock-in locus is AGCTCGCGTCTGCGCAGAG.

### RNA-seq and differential gene expression analysis

Progenies from heterozygous *dscaml1* mutant parents were anesthetized and harvested at 3.5-4 dpf. The anterior half of the animal was used for RNA preparation using the RNA Miniprep Kit (Zymo). The posterior half was used for genotyping. Three biological replicates for each group were analyzed, each containing RNA from 6-11 animals. All samples had RIN ≥ 8.0 and were converted into a strand-specific library using Illumina’s TruSeq Stranded mRNA HT Sample Prep Kit (RS-122-2103; Illumina) for subsequent cluster generation and sequencing on Illumina’s NextSeq 75 sequencer. Sequence data processing, alignment, read count, mapping, and quality control were performed as previously described (Ates et al., 2020). Differential expression was tested for significance using the false discovery rate (FDR) (Benjamini-Hochberg) corrected Likelihood Ratio Test (LRT) in the R-package DESeq2 (Love et al., 2014). 238 and 116 genes showed a significant difference in read counts at FDR<0.01 and 0.001, respectively. Original sequence data have been deposited in NCBI’s Gene Expression Omnibus (Edgar et al., 2002) and will be accessible through GEO Series accession number GSE213858.

### Fluorescent *in situ* hybridization and immunohistochemistry

Single and double whole-mount fluorescent *in situ* hybridization (FISH) was performed using protocols described previously (Pan et al., 2012). Probes were synthesized by *in vitro* transcription using the DIG and Fluorescein RNA Labeling Mix (Roche). DIG and fluorescein-labeled probes were detected with anti-DIG or anti-Fluorescein POD-conjugated Fab fragments (Roche) and Cy3 or Fluorescein TSA-plus Reagent (Akoya Biosciences). Plasmid template for *crhb* (Lohr et al., 2009) was provided by Dr. David Prober at Caltech. The *dscaml1* probe was generated as described previously (Ma et al., 2020b).

Immunohistochemistry was performed as described previously (Ma et al., 2020a). Nuclei were stained with TOTO-3 Iodide (ThermoFisher). RFP was stained with chicken anti-RFP (600-901-379S, Rockland) or rabbit anti-RFP (PM005, MBL Life Science). GFP was stained with rabbit anti-GFP (598, MBL Life Science). CRH was stained with rabbit anti-CRH (PBL rC68) provided by P. Sawchenko and J. Vaughan from the Salk Institute. All FISH and immunohistochemistry samples were mounted in 1.5% low-melt agarose in glass-bottomed Petri dishes (P50G-1.5-14-F; MatTek) and imaged using a Nikon A1 upright confocal microscope.

### Cortisol extraction and ELISA

A detailed protocol for cortisol extraction and ELISA is provided in the online supplementary method. At 4.5 dpf, *dscaml1-/-* and control animals were separated based on the darker pigmentation of the *dscaml1-/-* animal (Ma et al., 2020b). 30-35 animals were placed in each petri dish for cortisol extraction. The morning baseline (unstressed) sample collections were done at 15-30 minutes after light onset (08:15-08:45), and the afternoon sample collections were done between 14:30-16:00. Hyperosmotic stress and stirring stress experiments were done between 14:30-15:30. 6 biological duplicates—each containing a pool of 30 animals—were collected for each genotype (*dscaml1-/-* and control) and stress condition (morning baseline, afternoon baseline, stirring stress, osmotic stress). Mutants and control animals are tested side by side for each experiment.

Stirring stress was induced by creating a vortex water flow with a spinning magnetic stir bar (Castillo-Ramirez et al., 2019). A small magnetic stir bar was placed into a 100 mm petri dish containing 35 animals and 20 ml of E3 media. The stir bar was rotated at 300 rpm with a stirring microplate for 5 minutes. Hyperosmotic stress was induced by increasing salt concentration in the media (Yeh et al., 2013). 30 animals were placed in 8 ml of E3 media. Then, 2 ml of prewarmed 1.25M NaCl was added to the media for a final concentration of 250 mM for 20 minutes.

Sample homogenization and cortisol extraction were performed as described by Yeh et al. (Yeh et al., 2013). Briefly, 5 dpf larvae were rapidly immobilized with ice-cold E3 media and then flash-frozen at −80°C. Once all samples were collected, cortisol from the frozen samples was extracted with ethyl acetate (33211-1L-R; Sigma-Aldrich). Cortisol concentration was measured using a commercial ELISA kit, following the manufacturer’s instructions (500360; Cayman Chemical). Sample plates were read with a microplate reader (FilterMax F3; MicroDevices) 90-120 minutes after initial development.

### Live imaging of CRH neurons

Cre-mediated recombination of the *crhb:LRLG* transgene was induced by injecting Cre mRNA into the embryo at the 1-cell stage. To achieve partial Cre-mediated recombination, ~30 pg of CreER mRNA was injected at the 1-cell stage and 4-Hydroxytamoxifen (4-OHT) was added to the embryo media at 6 hpf (10 μM), followed by washout with E3 media at 24 hpf. To achieve complete Cre-mediated recombination, ~50 pg of *in vitro* transcribed Cre.zf1 mRNA (#61391, Addgene) was injected into the embryos (Horstick et al., 2015).

To perform confocal live imaging, 3 dpf animals were anesthetized with 0.01% tricaine methanesulfonate (MS-222, Sigma) and embedded in 1% low-melting point agarose, with the dorsal side resting on the glass surface inside the glass-bottomed petri dish (P50G-1.5-14-F; MatTek) (Beier et al., 2016). The petri dish was then filled with E3 media containing 0.01% tricaine methanesulfonate (MS-222, Sigma). Confocal z-stacks were acquired every 15 or 30 minutes for 12 hours on a Nikon A1 confocal microscope.

Two-photon live imaging was done on a custom Bruker two-channel two-photon microscope. A tuneable Ti:Saphire laser (Chameleon Vision II; Coherent) was tuned to 980 nm to excite RFP and GFP simultaneously. 78 hpf animals were anesthetized and embedded the same way as confocal live imaging, but with the dorsal side away from the cover glass. Each animal was imaged at 78, 84, 96, 108, and 120 hpf (Fig. 7A). After each time point, imaged larvae were gently removed from the agarose and recovered in E3 media at 28.5°C under normal day/night cycles. After the last time point, genomic DNA was prepared for all imaged animals and genotyped.

### Image Processing and Statistical Analyses

Images were processed using Fiji—an open-source image processing software (Schindelin et al., 2012). For FISH images, images were convolved (kernel=12) to enhance cell boundaries, and the center of each cell was manually tagged using the ROI manager tool. The cell number equals the number of ROIs in each animal. The signal intensity per cell was defined as the median signal intensity of all ROIs in a given animal. The number of RFP+ cells in *crhb:LRLG* animals were counted using the ROI manager tool without convolution. For confocal live imaging, images were pre-processed using the Denoise.AI function in the Nikon Elements software. For two-photon imaging, z-stacks from different time points were aligned using the “Correct 3D drift” function. Spectral overlap between the RFP and GFP channels was linearly unmixed. Individual cells were tracked with the MTrackJ plugin in Fiji (Meijering et al., 2012).

All statistical analyses were performed in GraphPad Prism (Version 9). For normally-distributed data, parametric tests (*t-*test or ANOVA) were used. For non-normally distributed data, non-parametric tests (Mann-Whitney) were used. The Holm-Sidak post-test was used to correct for multiple comparisons, and the adjusted *p* values are shown. All values are expressed as mean ± standard error, unless otherwise noted. Statistical tests were considered significant when *p*<0.05.

## AUTHOR CONTRIBUTIONS

M.M and Y.A.P conceived and designed the experiments. M.M and A.A.B prepared samples for RNA sequencing. Y.A.P analyzed the RNA sequencing data. M.M performed ELISA and histochemistry experiments and analyzed the data. M.M and K.C.C performed *in situ* hybridization experiments and analyzed the data with contributions from C.S, and K.S. S.H generated the *crhbl:LRLG* transgenic line. M.M and Y.A.P wrote the manuscript. Y.A.P provided project administration and acquired funding.

## ACKNOWLEDGEMENTS

We thank the animal care staff and veterinarians for animal husbandry; members of the Pan laboratory for helpful discussions; R. Settlage for computational analysis of RNA-seq data; S. Ryu and C. Yeh for sharing unpublished data; S. Imani for help with the quantification of IHC data; M. Wagle and S. Guo for advice on cortisol extraction procedures; M. Fox and A. Morozov for constructive feedback on the manuscript.

## DATA AVAILABILITY

Next-generation sequencing data utilized in this publication are available from the Gene Expression Omnibus (accession code GSE213858).

---

**Supplementary Fig. 1.**
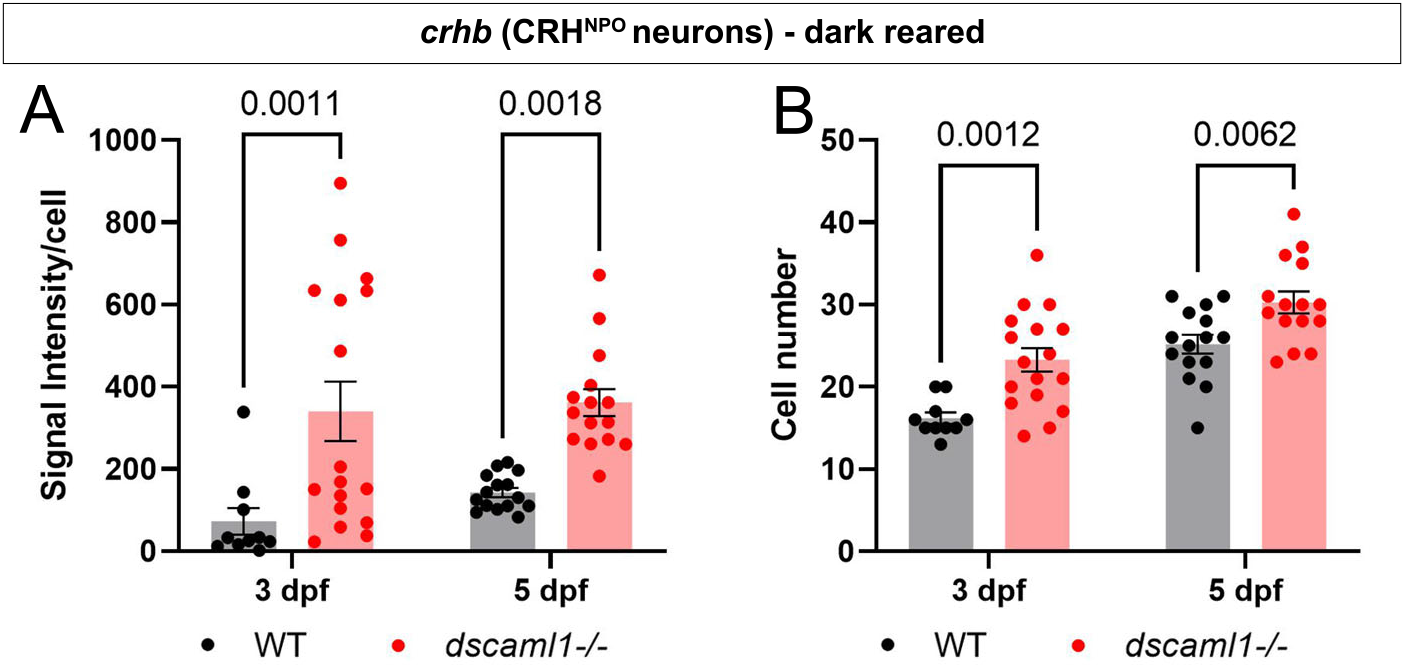
CRH^NPO^ neuron development is impaired in the absence of ambient light (dark reared). Quantification of CRH^NPO^ neurons, labeled by *crhb* FISH. Graphs show the signal intensity per cell (A) and cell number (B). Multiple-comparison corrected *p* values are as shown. WT: n=10 (3 dpf), 15 (5 dpf). *dscaml1-/-:* n=17 (3 dpf), 15 (5 dpf)

**Supplementary Fig. 2.**
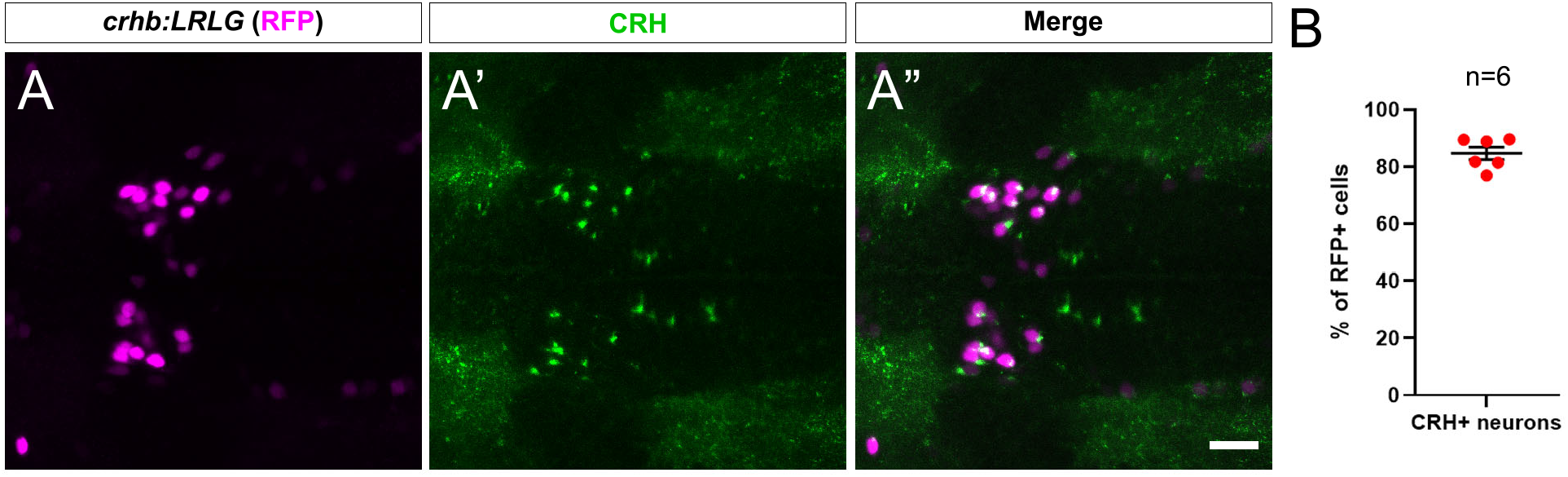
The *crhb:LRLG* transgenic line labels CRH-expressing neurons. (A-A”) Most RFP+ neurons (magenta, anti-RFP) express the CRH protein (green, anti-CRH). (B) Percentage of RFP+ neurons that express CRH. Mean, standard error, and sample size are shown. The scale bar is 20 μm.

**Supplementary Fig. 3.**
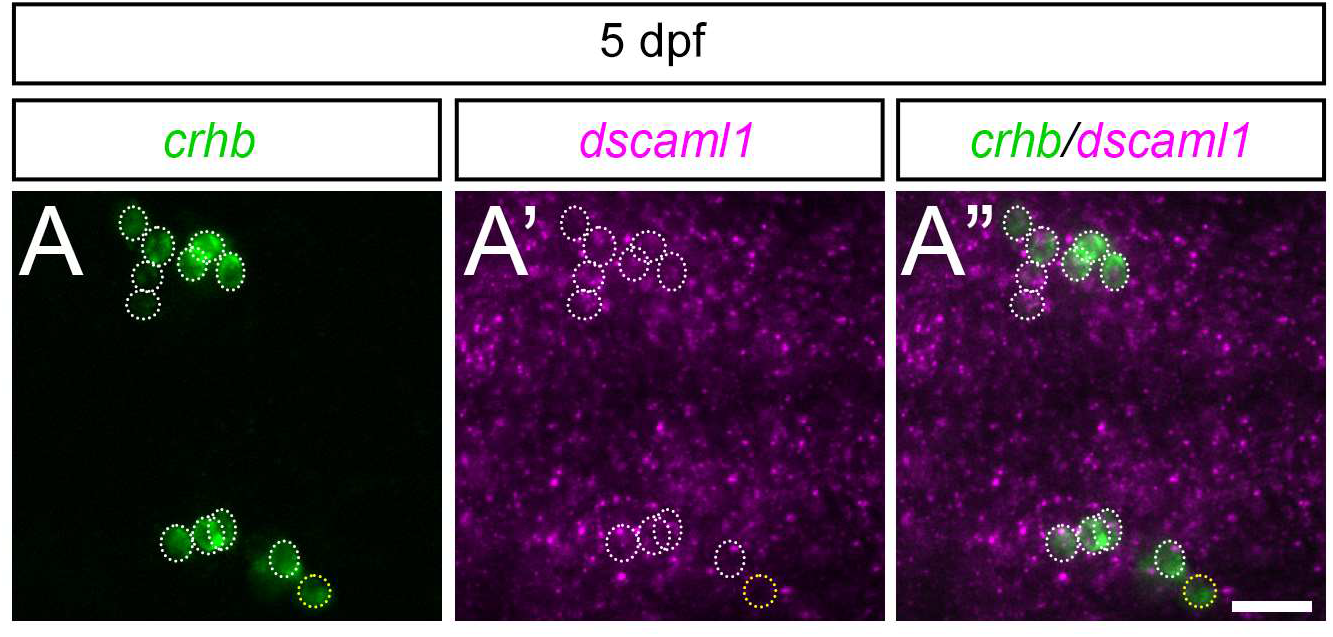
*dscaml1* is expressed in CRH^NPO^ neurons during larval development. Double FISH with *crhb* (green, A) and *dscaml1* (magenta, A”) RNA probes at 5 dpf. Merged image is shown in A”. Dotted circles mark the outline of *crhb+* cells that express *dscaml1* (white) or do not express *dscaml1* (yellow). The scale bar is 20 μm.

**Supplementary Video 1.** Time-lapse movie of *crhb:LRLG* larvae. CRH neurons were labeled with RFP (magenta) and GFP (green). Two cells (arrowheads, one green and one magenta), the same ones as shown in Fig. 5H-H”, move away over time.

**Supplementary Video 2.** Time-lapse movie of larvae with CRH neurons labeled with GFP (green) and microglia labeled with mCherry (magenta). The red “X” marks the cell indicated in Fig. 3I-J’. Maximum projection of z-stack for each time point is shown.

