## Supplementary material for "Deficiency in the cell-adhesion molecule *dscaml1* impairs hypothalamic CRH neuron development and perturbs normal neuroendocrine stress axis function": Supp methods

**Supplementary document, Ma et al. 2022.**

**Cortisol ELISA Assay using Cayman Cortisol ELISA Kit**

**ELISA kit**

Cayman Chemical #500360. Kit booklet: <https://cdn.caymanchem.com/cdn/insert/500360.pdf>

**Larvae raising**

After collecting eggs from breeding tanks, pre separate the eggs into 35 embryos per dish (use dishes with 60mm x 15mm, 10ml E3 media), change medium once at 2dpf, and make sure there are at least 30 live larvae per dish, and use 30 larvae after euthanized them with ice-cold E3. Keep fish on light background during raising. Segregate *dscaml1*-/- larvae by pigmentation if needed on 4-5dpf in the mornings while cortisol is high.

**Sample collection (Can be done over multiple days depending on exp design)**

1. Foremost, test all the tubes and pipette/pipette tips for compatibility with ethyl acetate before collecting fish into the tubes. Some pipettes/tubes could melt in ethyl acetate.
2. Prepare two buckets of ice, a bucket of ethanol/dry ice bath, and a bucket of dry ice.
3. Prepare ice-cold E3 in 50ml conical tubes by filling the conical tubes with E3, keep tubes in -20 for at least 15min before numbing the samples. Then keep the tubes on ice. The number of tubes depends on the number of dishes of fish.
4. Have a 6-well or multiple 6-well plate(s) ready on ice, fill each well half full with ice cold E3, and keep on ice, or could fill the wells 15min before experiment and place in -20, then move on ice when needed. Immerse the mesh-bottom inserts (24 mm Netwell™ Insert, Corning #3480) in the wells, and make sure it is permeabilizing properly.
5. Keep collection tubes on dry ice with caps labeled and opened, and arrange them in the order of the samples. (The tubes are hard to open and label once cold, so open and label the caps in advance.)
6. After stress treatments or when fish are ready to be numbed, pour ice-cold E3 into the dishes. Completely fill each dish with ice-cold E3. Then remove half of the volume with a glass transfer pipette without removing the fish, then pour more ice-cold E3 to fill the dish.
7. Pour larvae over the mesh insert, use a squirt bottle or glass transfer pipette to remove the larvae stuck on the dish into the mesh insert.
8. Move the mesh insert into the 6-well plate then fill up the dish with ice cold E3, making sure the mesh inserts are fully immersed. If fish are floating on the surface, lift the mesh insert up and down a few times.
9. Quickly pick 30 larvae with a glass transfer pipette into the pre-chilled 1.5ml tubes on dry ice, centrifuge the tubes for 5 sec and transfer the tubes on ice, remove excess E3, leave no more than ~25 ul along with larvae.
10. Flash freeze the samples in ethanol/dry-ice bath. Then store the tubes in -80°C until the day of ELISA process. Be aware of not touching the cap with ethanol, make sure the tube label is clear at all times.

**Cortisol extraction from larvae (Needs 1 full day)**

1. Prepare ice-cold PBS the same way when numbing the larvae.
2. Prepare duplicate tubes for BCA assay.
3. Thaw samples from -80°C on ice. If there are many samples, I try doing only two random tubes at a time until I add organic solvent, I keep the tubes on ice, then repeat the procedure for another two tubes to avoid degradation of cortisol after thawing.
4. Add 250 ul PBS to the tube after thawing, then homogenize the samples for 1min with a handheld homogenizer and pestle. Change out the pestle for each different tube.
5. Centrifuge the tubes for 5 sec.
6. Keep 50ul in a separate tube for total protein estimation or genotyping later, store at -20°C, use BCA microplate kit (if needed, else ignore this step).
7. Add 500 ul ethyl acetate (Sigma-Aldrich, 33211-1L-R) to each homogenate tube, vortex 30sec-1min at max speed, then leave on ice until all tubes are ready for high-speed centrifuge.
8. Separate solvent and aqueous phase by centrifuging the tubes for 5min at 3000 g – 5000 g, ideally at 4°C, if not available, room temperature is fine.
9. After phase separation, carefully transfer the organic solvent layer to a new 1.5ml tube with new pipette tips for each tube. Repeat previous steps (Steps 4-6) 1-2 more times to extract maximum cortisol. I do 500ul the first time, then 250ul for the next two times. Do not extract more than 1ml, or else it will spill out when vacuum concentrating with lids open.
10. Store aqueous layer for later in -80°C.
11. Evaporate organic solvent in the tubes by using a speed-vacuum concentrator for 30-40 min at medium heat (30-43°C work fine, depending on the choices provided by the machine). Extend the time if needed. Check each tube when done, make sure there is no liquid left in the tubes. Residual organic solvent could disrupt the ELISA reaction.
12. Reconstitute precipitant containing cortisol in 750ul-1000ul 1X ELISA buffer, prepare buffer according to Cortisol ELISA kit instructions. Cayman ELISA kit Page 12 “PRE_ASSAY PREPARATION”.
13. If using samples immediately, keep at 4°C with occasional vortexing for 14-24hrs, at 1200rpm, repeat several times in the hours. Then spin down for 5 sec. If samples are not used immediately, freeze at -20°C. Upon use, thaw and vortex 5min at 1200rpm. Spin down for 5 sec.
14. Use 50 ul of the reconstituted cortisol for ELISA following the ELISA kit. When making a second dilution, add the reconstituted cortisol to ELISA buffer at desired dilution (1:5, or 1:10), make sure to vortex thoroughly to mix, then spin down for 5 sec. Repeat expelling can’t mix cortisol and ELISA buffer thoroughly when ELISA buffer is at a low temperature or when too much cortisol is clogging the pipette tips.

**Performing ELISA with Cayman ELISA Kit Assay (Needs 1.5-2 full days)**

Follow cortisol ELISA kit protocol

Read the ELISA kit manual carefully and thoroughly to get familiar with the reagents and terms.

**Tips**

\# Make ELISA buffer (ELISA kit Page 12) at the last step of cortisol extraction. ELISA buffer will be used on the first day of ELISA assay. One bottle of ELISA buffer is enough for 1 full 96 well plate exp.

\# Make Wash buffer (ELISA kit Page 12) after the plate is at 4°C. 150-200 ml volume is enough for 1 full 96 well plate exp.

\# Save the adhesive cover from the first day ELISA overnight incubation, it will be needed again on the second day during plate development.

\# Make Ellman’s Reagent after plate washing.

\# Use new non-filter tips for removing reagents for each well during washing.

\# Use reagent reservoirs and repeat pipettors when washing.

\# Before developing the plate, use 200ul of wash buffer/time to rinse the wells 5-6 times.

**11-07-19 created by Michelle Ma, 5/17/22 edited by Albert Pan**
